# Robust activity-dependent mitochondrial calcium dynamics at the AIS is dispensable for action potential generation

**DOI:** 10.1101/2025.05.12.653506

**Authors:** Koen Kole, Maarten H.P. Kole

## Abstract

Mitochondria are diverse and multifaceted intracellular organelles regulating oxidative energy supply, lipid metabolism and calcium (Ca^2+^) signaling. In neurons the spatial sequestration of cytoplasmic Ca^2+^ by mitochondria plays a critical role in determining activity-dependent spine plasticity, shaping the presynaptic transmitter release characteristics and contributing to sustained action potential firing. Here, we tested the hypothesis that mitochondria at the axon initial segment (AIS) affect the microdomain cytoplasmic Ca^2+^ transients, thereby regulating Ca^2+^-dependent voltage-gated ion channels at the plasma membrane and initiation of action potentials. Using 3D electron microscopy (EM) reconstructions and virally injecting genetically encoded fluorescence indicators we visualized the ultrastructure and distribution of mitochondria selectively in thick-tufted layer 5 pyramidal neurons. We found that most mitochondria were stably clustered to the proximal AIS, while few were observed at distal sites. Simultaneous two-photon imaging of action potential-dependent cytoplasmic and mitochondrial Ca^2+^, combined with electrophysiological recordings showed the AIS mitochondria exhibit powerful activity-dependent cytosolic Ca^2+^ uptake. However, while intracellular application of the mitochondrial Ca^2+^ uniporter inhibitor Ru360 fully blocked mitochondrial Ca^2+^ import, it did not affect action potential input-output function, action potential dynamics nor the ability to produce high-frequency burst output. Together, the results indicate that AIS mitochondria are dispensable for temporal and rate encoding, suggesting that mt-Ca^2+^ buffering at the AIS may be involved in non-electrical roles, including AIS maintenance or axonal transport.

## Introduction

Mitochondria are organelles of the eukaryotic cell fundamentally tasked with the production of the cellular energy carrier adenosine triphosphate (ATP). Emerging insights have shown additional and versatile functions of mitochondria including lipid synthesis, regulation of cell death and, notably, calcium (Ca^2+^) homeostasis (Rizzuto *et al*., 2012). The multifaceted functionality of mitochondria is particularly clear in neurons, which are highly dependent on both ATP and Ca^2+^ homeostasis to enable the faithful propagation of action potentials and synaptic vesicle release (Attwell & Laughlin, 2001; Burgoyne, 2007; Hallermann *et al*., 2012). In neurons, the strong morphological polarization and compartmentalization of the cytoarchitecture is also reflected by the diversity in the mitochondrial pool. For instance, mitochondria in the dendrites are elongated and support spine plasticity (Lewis *et al*., 2018; Rangaraju *et al*., 2019) whereas axonal mitochondria are short and sustain synaptic vesicle release (Ashrafi *et al*., 2017, 2020). Beyond ATP synthesis, the mitochondrial Ca^2+^ (mt-Ca^2+^) buffering within axons plays and important role in local Ca^2+^ dependent processes. Within presynaptic terminals mitochondria take up Ca^2+^ and sequester it, lowering local cytosolic Ca^2+^ levels and reducing synaptic vesicle release rates (Kwon *et al*., 2016; Lewis *et al*., 2018). Mt-Ca^2+^ buffering is thus an effective means by which neurons can regulate their cytosolic Ca^2+^ levels within subcompartments, allowing high spatial and temporal control over Ca^2+^-dependent pathways.

Interestingly, by Ca^2+^ buffering mitochondria also impact the electrophysiological membrane properties of neurons. Cytosolic Ca^2+^ entering the mitochondrial matrix in part drives ATP production via oxidative phosphorylation (Rizzuto *et al*., 2012; Szibor *et al*., 2020), thereby supplying energy to for example the sodium (Na^+^)/potassium (K^+^) ATPase pump (Stoler *et al*., 2022; Zampese *et al*., 2022). Such fueling of the Na^+^/K^+^ ATPase restores ionic concentrations and sustains spiking rates (Lin *et al*., 2019; Stoler *et al*., 2022; Zampese *et al*., 2022). Previous studies in cortical pyramidal or vasopressin neurons of the hypothalamus show that mitochondria-dependent regulation of the membrane excitability adjusts the firing rate as well as the Ca^2+^-dependent slow afterhyperpolarization current (Styr *et al*., 2019; Groten & MacVicar, 2022; Kirchner *et al*., 2024). It remains to be tested if Ca^2+^ buffering by mitochondria also impacts other electrophysiological properties, such as firing frequency, bursting or action potential waveform. Moreover, it is unknown whether the previous observations result from the collective Ca^2+^ buffering capability of all mitochondria in a given neuron, or if there is a specific locus where mt-Ca^2+^ buffering controls neuronal electrophysiological properties. One highly specialized subcompartment of the neuron is the axonal initial segment (AIS), where action potentials (APs) are initiated by the clustering of high densities of voltage-dependent ion channels (Kole & Stuart, 2012; Jenkins & Bender, 2024). While the membrane excitability at the AIS axolemma is dominated by the voltage-dependent Na^+^ and K^+^ channels, AP generation is also shaped by voltage-dependent Ca^2+^ influx and cytoplasmic Ca^2+^ concentrations. For example, blocking voltage-gated Ca^2+^ channels locally at the AIS increases the AP threshold and delays AP initiation within high-frequency bursts (Bender & Trussell, 2009). Furthermore, Ca^2+^ not only charges the membrane but also acts as a downstream signalling molecule: axonal Ca^2+^-activated K^+^ (K_Ca_) channels regulate AP decay time and decrease neuronal excitability (Yu *et al*., 2010; Gründemann & Clark, 2015; Filipis *et al*., 2023). Several voltage-gated ion channels possess Ca^2+^-calmodulin sensing domains, mediating Ca^2+^-dependent regulation of kinetics of voltage-gated sodium channels (Ben-Johny *et al*., 2014; Wang *et al*., 2014) or closure of the Kv7 channel (Delmas & Brown, 2005; Martinello *et al*., 2015; Bernardo-Seisdedos *et al*., 2018). The Ca^2+^ dependent regulation of ion channels require neurons to carefully control their cytoplasmic Ca^2+^ levels at the AIS, which can be achieved in part by sequestering it internally and regulating its release via stores (Verkhratsky & Petersen, 1998).

Mitochondria have been reported in the AIS, but their function in this domain remains poorly understood (Dimova & Markov, 1976; Li *et al*., 2004; Kole *et al*., 2022; Tjiang & Zempel, 2022). The main entry route for Ca^2+^ into the mitochondrial matrix is via the mitochondrial calcium uniporter (MCU), a selective Ca^2+^ channel residing in the mitochondria inner membrane. The MCU has a low Ca^2+^ affinity, activated with cytoplasmic Ca^2+^ concentrations as low as ∼1 µM (Payne *et al*., 2017). AIS mitochondria in neocortical layer 5 (L5) pyramidal neurons may rapidly encounter such cytosolic Ca^2+^ levels since even a single AP raises cytoplasmic Ca^2+^ by ∼1 µM (Hanemaaijer *et al*., 2020). Here, we tested the hypothesis that mitochondrial Ca^2+^ buffering at the AIS regulates AP initiation and generation. We examined the anatomical distribution of L5 pyramidal neuron mitochondria by using 3D electron microscopy (EM) reconstructions and functionally studied their role by using genetically encoded tools to visualize mitochondria and their Ca^2+^ uptake, combined with electrophysiological recordings and pharmacological manipulation. We found that the AIS contains a gradient of mitochondria which are stably located and exhibit strong Ca^2+^ buffering. Intracellular application of the MCU pore complex inhibitor RU360 however did not alter AP firing properties. Our results reveal a striking axonal compartmentalization of mitochondrial distribution and Ca^2+^ buffering and suggest that mt-Ca^2+^ buffering at the AIS is dispensable for AP generation in L5 pyramidal neurons.

## Results

### Mitochondria densely populate the AIS

To examine the distribution of mitochondria in L5 pyramidal neuron AISs at ultrastructural detail we made use of a publicly available annotated 3D EM dataset of the adult mouse primary visual cortex (Consortium *et al*., 2025). We randomly selected thick-tufted L5 pyramidal neurons and segmented mitochondria at the AIS (excluding the first 3 µm representing the axon hillock, the transition from soma into the AIS) and the first myelinated internode, quantifying their occupancy, size and density (**Figure 1A-C**). At the AIS, defined as the unmyelinated region between the axon hillock and first paranodal loops, mitochondria occupied mostly the proximal AIS and significantly reduced in probability at the distal 10 µm, in line with previous reports (Tamada *et al*., 2021; Tjiang & Zempel, 2022) (**Figure 1Di**). Within internodes, mitochondria tended to avoid the first ∼2 µm representing the paranode but were otherwise uniformly distributed (**Figure 1Dii**). We observed that ∼75% of first nodes of Ranvier (branch points after the first internode) harbored mitochondria (**Figure 1Dii**). Considering the AIS as a whole, mitochondria were present at significantly higher densities compared to the first internode (0.67 ± 0.24 vs. 0.38 ± 0.10 mitochondria/µm; paired *t*-test P = 0.0089) and showed a trend to be smaller (AIS, 0.11 ± 0.05; Internode, 0.14 ± 0.06 µm^3^; paired *t*-test P = 0.0822). Given the declining mitochondrial occupancy in the distal AIS, we quantified the density and size of mitochondria in the first and second 50% of the total AIS length (proximal and distal AIS, respectively). The data showed that mitochondria were significantly more abundant within the proximal AIS at over twice the density (**Figure 1E**; prox AIS, 1.14 ± 0.18; dist AIS, 0.45 ± 0.10; internode, 0.38 ±0.04 mitochondria/µm; repeated one-way ANOVA P = 0.0011). In addition, mitochondria within the distal AIS were significantly smaller than in the internode (**Figure 1F**; prox AIS, 1.12 ± 0.02; dist AIS, 0.05 ± 0.02; internode, 0.15 ± 0.02 µm^3^; repeated one-way ANOVA P = 0.0132). This also tended to be true for the mitochondrial volume relative to that of the axon, but this did not reach statistical significance in *post hoc* tests (**Figure 1G**; prox AIS, 11.91 ± 1.69; dist AIS, 6.80 ± 2.28; internode, 12.61 ± 0.94 v/v; repeated one-way ANOVA *P* = 0.0369). Together, these results indicate that mitochondria densely populate the proximal AIS.

**Figure 1.**
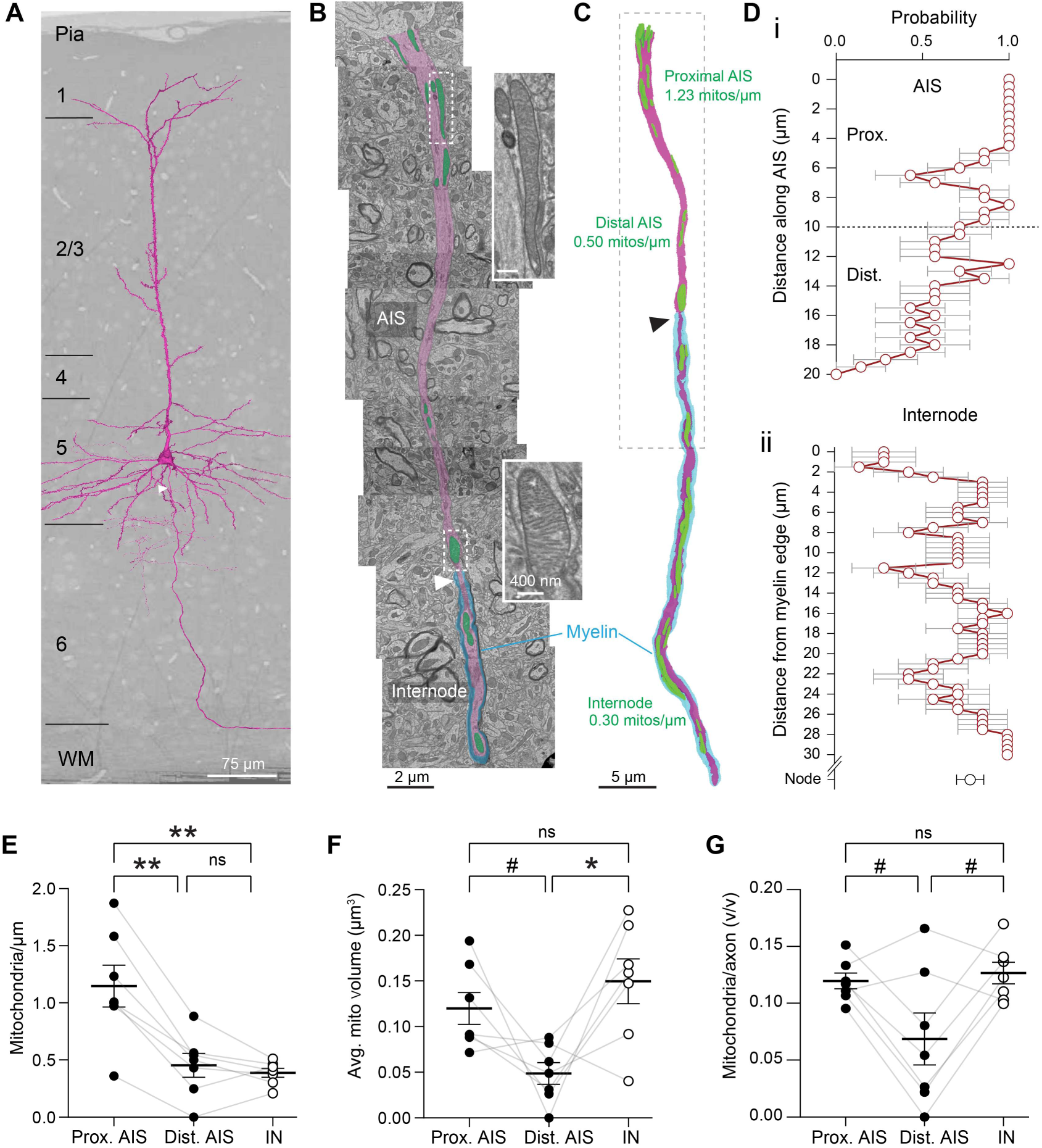
Mitochondria densely populate the proximal AIS. **A**. 3D reconstructed morphology of a thick-tufted layer 5 pyramidal neuron (magenta) projected on the EM block of a mouse visual cortex. White arrow, location of the axon. **B** Collage of single EM images of the AIS and a section of the first internode of the cell shown in A. AIS is pseudocolored in magenta, mitochondria are pseudocolored in green, myelin is pseudocolored in cyan. Note that this is a single plane, underestimating the number of mitochondria. Dashed boxes correspond to example mitochondria. White arrowhead, paranodal loops of first internode; **C** 3D reconstruction of the mitochondria and axon. Dashed box corresponds with the view in A, black arrowhead indicates paranodal loops of the first internode. **D** Distribution of mitochondria within the AIS (i) or internodes and nodes (ii), dashed line at 10 µm separating proximal versus distal AIS. Y axes are limited to lengths where each AIS/internode is represented; **E-G** Mitochondria are more densely populated at the proximal AIS compared to the distal AIS and first internode (**E**; repeated measures one-way ANOVA P = 0.0011; Tukey post-hoc test prox vs. dist AIS **P = 0.0011; Prox AIS vs. internode P = **0.0071; Dist AIS vs internode P = 0.6973). At the distal AIS, a trend of smaller mitochondria is observed both in absolute (**F**; repeated measures one-way ANOVA P = 0.0132; Tukey post-hoc test prox vs. dist AIS P = 0.0828; Prox AIS vs. internode P = 0.3109; Dist AIS vs internode P = 0.0419) and relative volume (**G**; repeated measures one-way ANOVA P = 0.0369; Tukey post-hoc test prox vs. dist AIS P = 0.0963; Prox AIS vs. internode P = 0.3434; Dist AIS vs internode P = 0.0797). D-G, n = 7 axons from 1 mouse.

To test further the distribution of mitochondria in molecularly identified AISs we next studied acute slices using previously developed Cre-dependent viral tools to fluorescently label mitochondria (Kole *et al*., 2022). *Rbp4*-Cre mice, which express Cre recombinase selectively in L5 pyramidal neurons (Gerfen *et al*., 2013), were injected in the primary somatosensory cortex with an AAV vector to express Cre-dependent mitochondria-targeted green fluorescent protein (mt-GFP). Immunofluorescence staining showed that this viral approach effectively and specifically labels mitochondria in L5 pyramidal neurons (**Figure 2A**). We then applied immunolabeling of the AIS-associated cytoskeletal protein ßIV-spectrin, to identify mitochondria within the molecularly identified AIS (**Figure 2A-C**). In acute brain slices, we made patch-clamp recordings from L5 pyramidal neuron to fill the cytoplasm with biocytin, allowing detailed reconstruction of mitochondria labeled mt-GFP expressing (mt-GFP^+^) at the AIS and first internode. In accord with the 3D EM data, the AIS contained mitochondria with significantly higher density in the proximal AIS, and mitochondria numbers declining with distance from the soma (**Figure 2C,D**; prox AIS, 0.45 ± 0.06; dist AIS, 0.23 ± 0.04; internode, 0.22 ± 0.01 mitochondria/µm; repeated one-way ANOVA P = 0.0033). Compared to 3D EM our immunohistochemistry data indicated lower mitochondrial densities (on average 0.36 vs. 0.67 mitochondria/µm; unpaired t-test, P = 0.0118). This is likely explained by the much higher spatial resolution that EM offers (4 nm), allowing the identification of smaller mitochondria that are below the detection threshold of confocal microscopy (> 400 nm).

**Figure 2.**
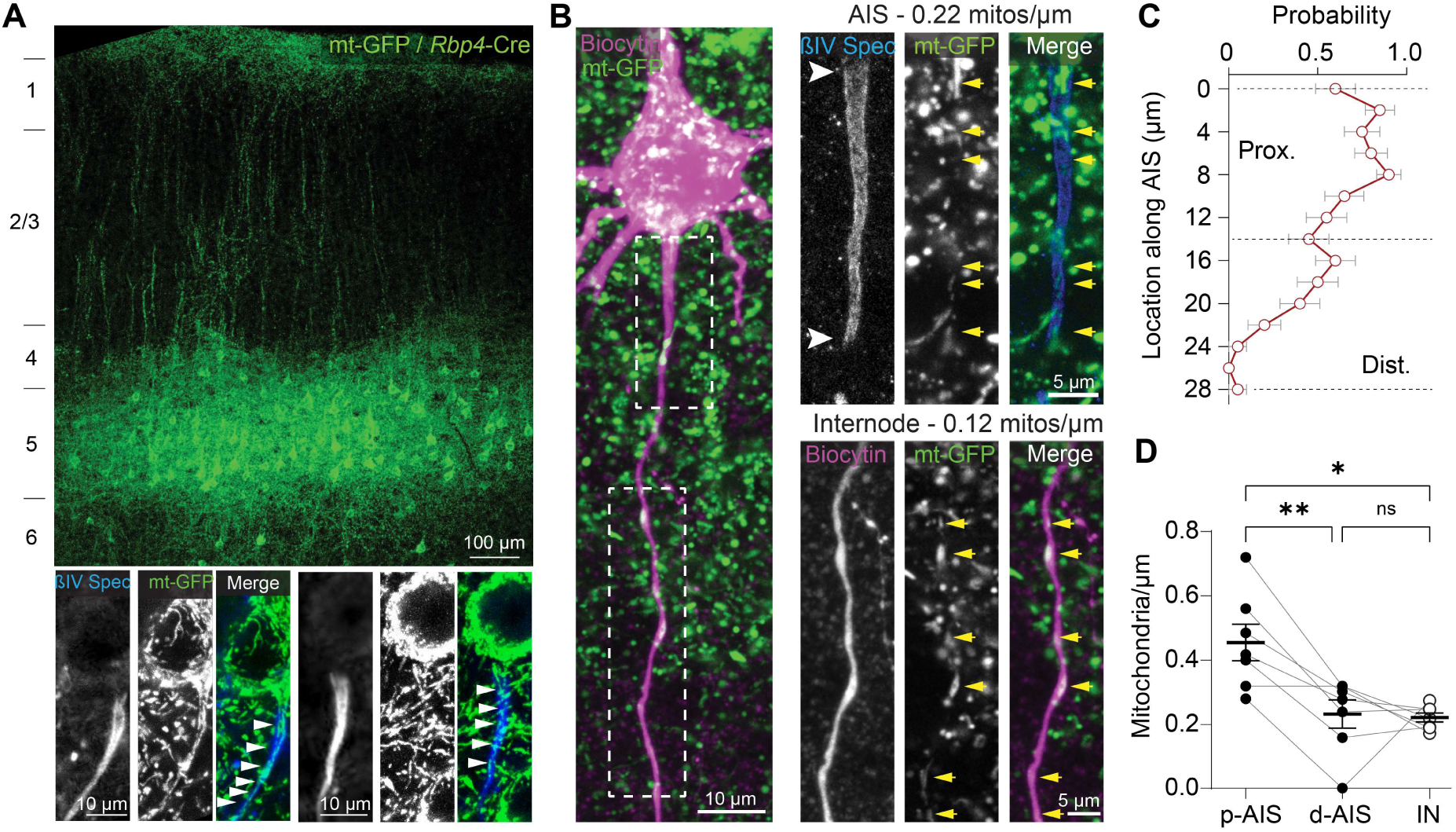
Mitochondria densely populate the proximal AIS. **A** Expression of mt-GFP in *Rbp4*-Cre mice labels mitochondria selectively in L5 pyramidal neurons. Insets show dense mitochondria at the AIS, visualized by β4 spectrin immunolabeling; **B** Example of a biocytin-filled mitoGFP^+^ L5 pyramidal neuron. Dashed boxes correspond to the high magnification images on the right (AIS and first internode). White arrowheads indicate AIS, yellow arrows indicate mitochondria; **C** Mitochondria are most prevalent at the proximal AIS and reduce in density along the AIS length; **D** Mitochondrial density is highest at the proximal AIS (average 0.36 vs 0.22 mitochondria/µm; repeated measures one-way ANOVA P = 0.0033; Tukey post-hoc test prox vs. dist AIS \*\**P* = 0.0097; Prox AIS vs. internode \**P* = 0.0282; Dist AIS vs internode *P* = 0.9783); C, n = 20 AISs (not biocytin-filled) from 2 mice; D, n = 7 cells from 5 mice.

We next asked whether the high mitochondrial density at the proximal AIS represented a stable population or if they were mitochondria merely transported via microtubules within the AIS antero- or retrogradely between the soma and the distal axon (Misgeld & Schwarz, 2017). We therefore made live two-photon recordings of mitochondria at the AIS in acutely prepared brain slices of mitoGFP-expressing mice (**Figure 3A**). The AIS was readily recognizable by its high mitochondrial content, negating the necessity of a cytosolic marker to identify the axon. Although some mitochondria were motile, we found that the majority (73.64 ± 21.43%) was stably present at the AIS for the duration of the imaging session (8 to 10 minutes, **Figure 3B**), in agreement with previous reports (Tjiang & Zempel, 2022). Motile mitochondria predominantly moved in an anterograde direction (**Figure 3C**; 89.35 ± 18.53%) and were significantly smaller compared to stable mitochondria (**Figure 3D**; nested t-test, *P* < 0.0001; 0.36 ± 0.21 vs. 1.46 ± 0.89 µm^2^). We found no correlation between size and speed (**Figure 3E**; Pearson *r* = –0.08265, *P* = 0.6881). Taken together, our findings from EM, immunohistochemistry and live imaging indicate that mitochondria localize mostly to the proximal AIS and are stable.

**Figure 3.**
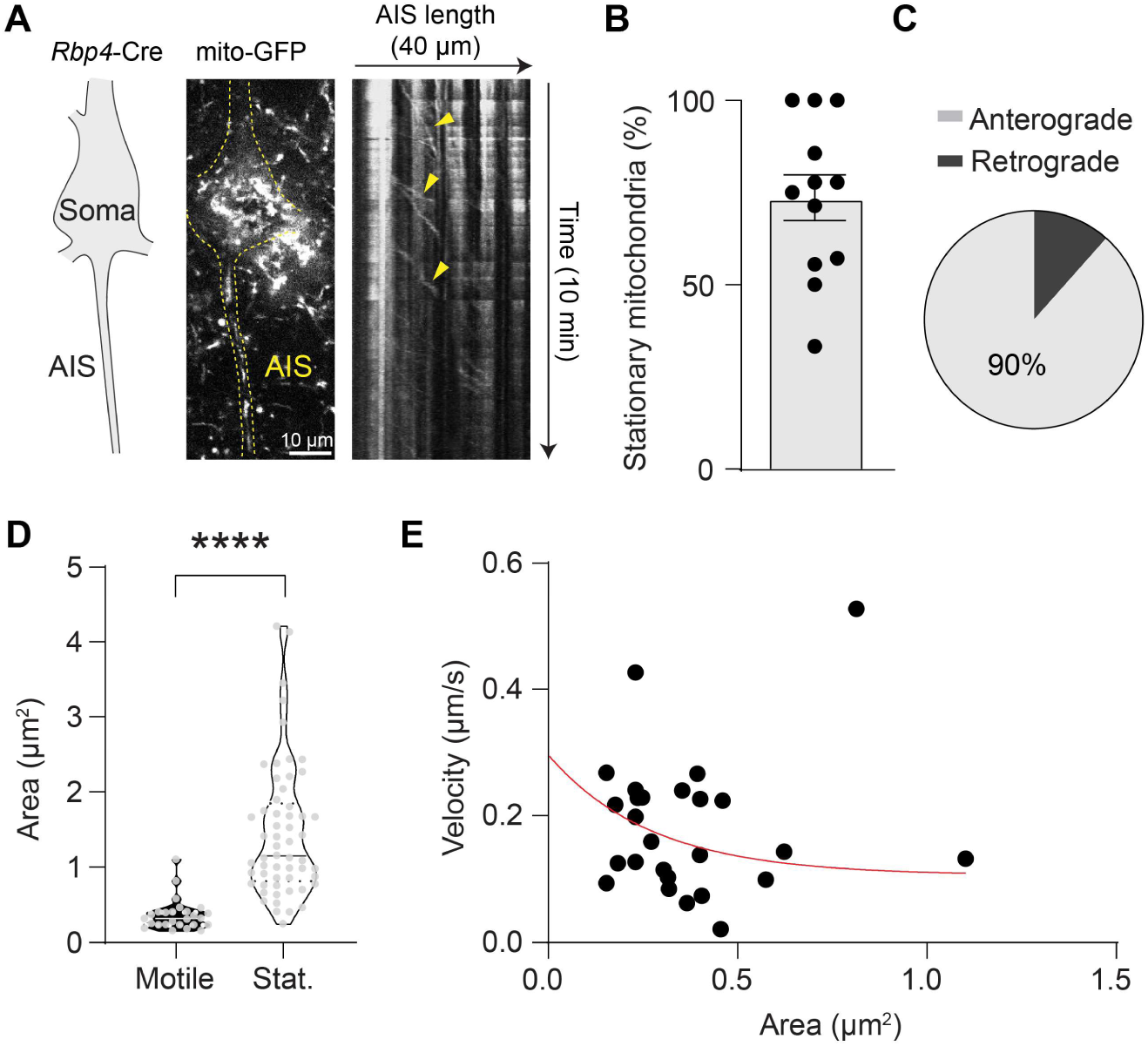
Stable and motile mitochondria at the AIS. **A** Example kymograph showing motility of some mitochondria (yellow arrows) but with most mitochondria remaining stably present during the imaging session; **B** Quantification of stationary mitochondria at the AIS; **C** Quantification of anterograde and retrograde movement by mitochondria at the AIS; **D** Stationary mitochondria are larger compared to motile mitochondria (nested t-test, \*\*\*\**P* < 0.0001); **E** No correlation between mitochondrial surface area and velocity (Pearson *r* = –0.08265, *P =* 0.6881). Red line indicates an exponential decay (*y* = 0.1871 × exp(–3.645 × *x*) + 0.1086). In B, each datapoint represents one AIS (n = 12 AISs from 5 mice); in D and E, each datapoint represents one mitochondrion (n = 27 mitochondria from 5 AISs from 5 mice).

### Mitochondria at the AIS buffer Ca^2+^

The AIS and nodes of Ranvier of layer 5 pyramidal neurons are characterized by large activity-dependent Ca^2+^ fluxes (Hanemaaijer *et al*., 2020). We therefore asked whether the activity-dependent Ca^2+^ buffering of mitochondria at the AIS differed from those in nodes, internodes and somata and how it correlated with cytosolic Ca^2+^ influx. To this end we expressed the genetically encoded, GFP-based and mitochondrion-targeted Ca^2+^ indicator mt-GCaMP6f selectively in L5 pyramidal neurons (Kole *et al*., 2022). We then performed whole-cell patch-clamp recordings in mt-GCaMP6f-expressing (Kd ∼375 nM; Chen et al.) L5 pyramidal neurons were recorded with intracellular solution supplemented with the red Ca^2+^ indicator dye Cal-590 (Kd 561 nM) (**Figure 4A**). The dual Ca^2+^ indicator approach enabled us to image the activity-dependent Ca^2+^ responses in the soma, AIS, myelinated internodes and nodes simultaneously in the cytosol (cyt-Ca^2+^) and mitochondria (mt-Ca^2+^). The data showed that during spike trains, mt-Ca^2+^ transients temporally followed those in the cytosol (**Figure 4B**). Indeed, in line with previous reports (Groten & MacVicar, 2022; Stoler *et al*., 2022), we found that mt-Ca^2+^ increase was significantly delayed by ∼150 ms (time to 50% peak; mt-Ca^2+^, 315.39 ± 18.37 ms, cyt-Ca^2+^, 149.43 ± 8.19 ms, unpaired t-test P < 0.0001; n = 9 cells from 6 mice) and showed a trend for slower decay time constant (mt-Ca^2+^, 426.20 ± 36.3 ms; cyt-Ca^2+^ 310.02 ± 57.15 ms, Mann-Whitney test P = 0.0770, n = 9 cells from 6 mice). It should be noted that we underestimate the decay duration, as our imaging sessions only captured the first few seconds of decay which in mitochondria can last for tens of seconds (Groten & MacVicar, 2022; Stoler *et al*., 2022).

**Figure 4.**
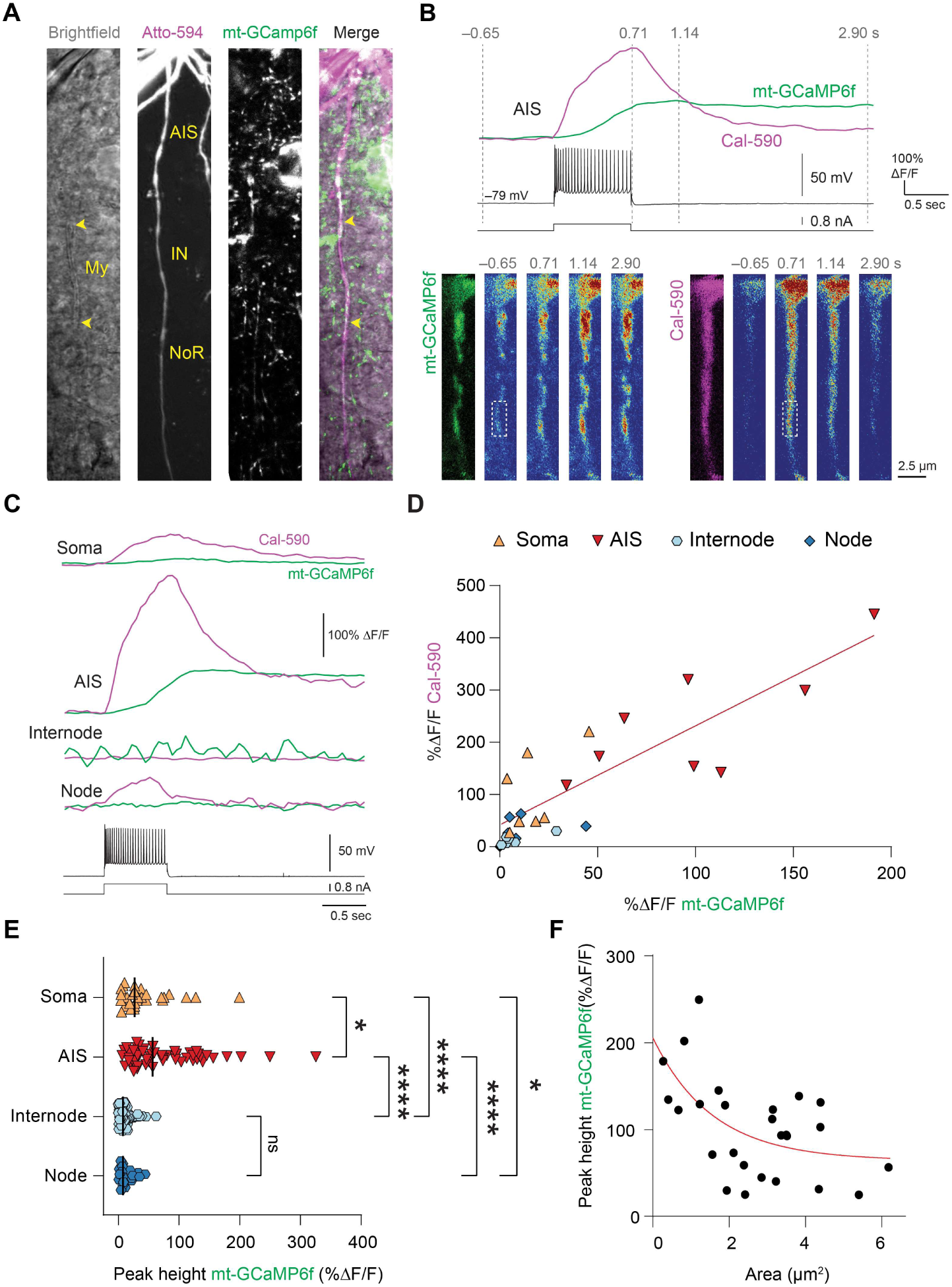
Activity-dependent cyt-Ca^2+^ and mt-Ca^2+^ dynamics at the AIS. **A** Example image of a mt-GCaMP6f^+^ L5 pyramidal neuron filled with Atto594 revealing the cytoarchitecture while transmission light reveals myelinated internodes (My, yellow arrowheads); **B** Top: Example traces of mitochondrial (mt-GCaMP6f, green) and cytosolic (Cal-590, magenta) Ca^2+^ transients in response to a 700 ms AP train (25 APs). Bottom: heatmaps of the corresponding traces at the indicated timepoints. Note the difference in onset and duration between mt- and cyt-Ca^2+^ responses. Dashed boxes indicate ROIs corresponding to traces. Scale bar represents 2.5 µm; **C** Example traces from different cellular compartments from the same cell of mt- and cyt-Ca^2+^ responses during a 700 ms AP train; **D** Scatter plot of mt- and cyt-Ca^2+^ responses at soma, AIS, internode and nodes. The strongest responses are observed at the AIS. Dashed line indicates a linear curve fit (*y* = 0.4096 + 1.896 × *x*); **E** Quantification of subcellular mt-Ca^2+^ responses. Datapoints indicate somata or, in the case of AIS, internode and node, individual mitochondria (Kruskal-Wallis test with Dunn’s *post hoc* test; AIS vs Soma \**P* = 0.0481, AIS vs Internode \*\*\*\**P* < 0.0001, AIS vs Node \*\*\*\**P* < 0.0001, Soma vs Internode \*\*\*\**P* < 0.0001, Soma vs. Node \**P* = 0.0417, Internode vs. Node *P* > 0.9999); **F** Smaller mitochondria exhibit stronger Ca^2+^ responses. Red line indicates a single exponential fit (*y* = 1.4145 × exp(–0.6339 × *x*) + 0.6435). D, n = 21 cells from 12 mice; E, n = 30 somata, 55 (AIS), 83 (internode) and 19 (node) mitochondria from 40 cells of 21 mice; F, n = 26 mitochondria from 8 AISs of 6 mice.

We then compared cyt-Ca^2+^ and mt-Ca^2+^ response amplitudes from the axonal subcompartments, which revealed that both cyt-Ca^2+^ and mt-Ca^2+^ responses were strongest in the AIS whereas they were very weak to nonexistent in myelinated internodes (**Figure 4B-E**), in agreement with previous observations (Hanemaaijer *et al*., 2020; Kole *et al*., 2022). In nodes we detected AP-dependent cyt-Ca^2+^ increase but these were irregularly paired with mt-Ca^2+^ transients (**Figure 4C-D**), suggesting that the Ca^2+^ influx may be insufficient for MCU activation. Given the variance in mitochondrial size (**Figure 1C**), we asked whether mt-Ca^2+^ responses varied by size and correlated mt-Ca^2+^ peak responses with surface area. This revealed that smaller mitochondria displayed larger Ca^2+^ transients (**Figure 4F**). We did not observe a correlation between mt-Ca^2+^ transient amplitude and distance from soma (Pearson correlation coefficient R^2^ = 0.0209, P = 0.4811). Taken together, these findings indicate that in L5 pyramidal neuron axons, mitochondria at the AIS strongly buffer activity-dependent Ca^2+^.

### Mt-Ca^2+^ buffering does not affect electrophysiological properties in L5 pyramidal neurons

Mt-Ca^2+^ buffering has previously been shown to attenuate cyt-Ca^2+^ concentration and shorten the duration of the slow afterhyperpolarization (AHP) (Groten & MacVicar, 2022). Since cyt-Ca^2+^ at the AIS of L5 pyramidal neuron reaches high concentrations, shaping the AP and slow AHP (Hanemaaijer *et al*., 2020; Roshchin *et al*., 2020) we hypothesized that mt-Ca^2+^ buffering in L5 pyramidal neurons affects the membrane excitability. To test this, we performed whole-cell patch clamp recordings while infusing the selective MCU inhibitor Ru360, which blocks the MCU by binding it at the mitochondrial intermembrane space (Baughman *et al*., 2011), effectively abolishing mitochondrial Ca^2+^ uptake. We observed that 45 minutes of Ru360 infusion attenuated mt-Ca^2+^ with an average reduction of 81% compared to control cells (**Figure 5A-D**; control, 2.74 ± 2.06; Ru360, 0.51 ± 0.30 ΔF/F, *P* = 0.0459, unpaired t-test). Next, we examined the role of blocking mt-Ca^2+^ on membrane excitability. By injecting temporally precise brief currents we evoked 100 APs at 100 Hz and quantified the slow AHP peak amplitude, halfwidth and area under the curve. Unexpectedly, and in contrast to previous work (Groten & MacVicar, 2022), we did not detect any effect of Ru360 on the AHP properties (**Figure 5E-H**; unpaired t-tests; peak *P* = 0.3228, halfwidth, *P* = 0.7871, AUC *P* = 0.4464). Although the slow AHP often takes >20 seconds to return to the resting membrane potential (Gulledge *et al*., 2013) also when we quantified the amplitude of the AHP at 2 seconds after AP stimulus termination there was no treatment effect detectable (unpaired t-test, *P* = 0.6313; membrane potential relative to rest: control, −1.86 ± 0.16; Ru360, −1.73 ± 0.17 mV). Furthermore, slower membrane currents beyond the recording window were unlikely to be affected since the resting membrane potentials were not affected by Ru360 (unpaired t-test, *P* = 0.8599; control, −77.42 ± 2.67; Ru360, −77.14 ± 3.23 mV).

**Figure 5.**
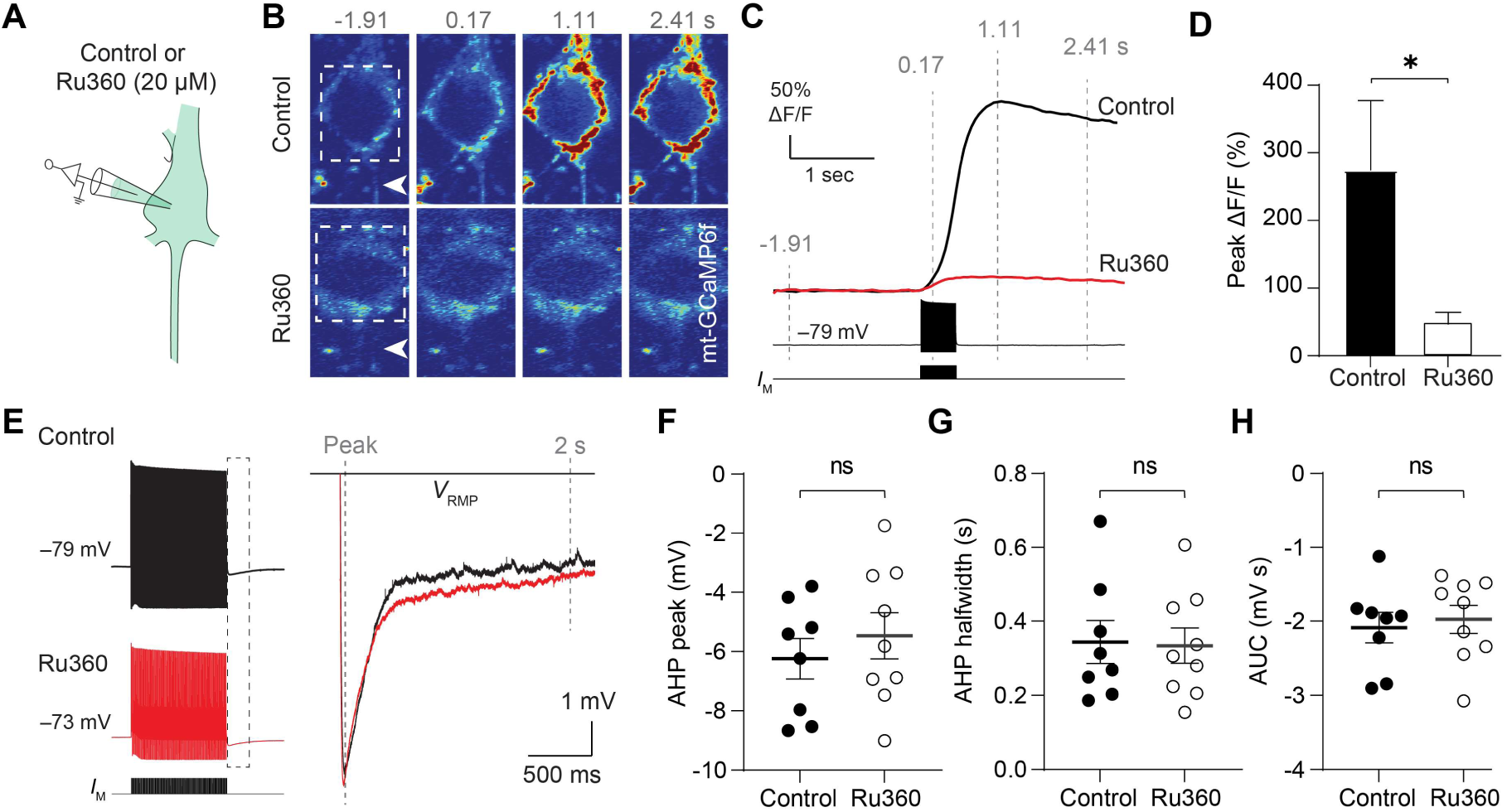
Blocking mt-Ca^2+^ buffering does not affect the slow AHP. **A** Schematic representation of the experiment. Whole-cell recordings are performed while infusing intracellular solution with or without Ru360. **B** Example mt-Ca^2+^ responses in a cell infused with intracellular without control (top) or with Ru360 (bottom). Arrowhead indicates proximal AIS; dashed box indicates the ROI used for the traces in C; **C** Traces corresponding to the example cells in B; **D** Quantification of somatic mt-Ca^2+^ responses in control or Ru360-infused cells (unpaired t-test, \**P* = 0.0459); **E** Example voltage traces (100 APs, 100 Hz) of a control or Ru360-infused cell. Dashed box corresponds to the inset on the right showing the AHP; **F-H** Quantification of the AHP peak (F; unpaired t-test, *P* = 0.3228), halfwidth (G; unpaired t-test *P* = 0.7871) and area under the curve (H; unpaired t-test, *P* = 0.4464). D: control, n = 4 cells from 2 mice; Ru360, n = 5 cells from 2 mice; F-H: control, n = 8 cells from 4 mice; Ru360, n = 9 cells from 5 mice.

While the slow AHP in L5 pyramidal neurons was independent from mt-Ca^2+^ buffering we postulated that other calcium sensing voltage-gated channels at the soma and AIS could be dependent on mt-Ca^2+^. The slowly activating Kv7 channels, expressed at high densities in the distal AIS, sets the neuronal resting membrane properties, regulates AP threshold and is gated by cytoplasmic Ca^2+^ (Delmas & Brown, 2005; Battefeld *et al*., 2014; Martinello *et al*., 2015; Bernardo-Seisdedos *et al*., 2018). With the Ru360-mediated loss of mt-Ca^2+^ buffering (**Figure 5**) the putatively increased cytoplasmic Ca^2+^ is expected to produce a closure of the Kv7 channel and thereby increase the neuronal excitability, by lowering the current threshold for AP generation. To explore this possibility, we injected incremental current steps and compared AP generation in control and Ru360 treated cells. Unexpectedly, we found no difference between the number of APs generated upon current injection (**Figure 6A-B**; two-way ANOVA; treatment effect *P* = 0.8250; interaction effect *P* = 0.8142;), neither a change of the rheobase current (unpaired t-test, *P* = 0.3355; control, 244.4 ± 84.57; Ru360, vs. 281.3 ± 65.12 pA) nor a change of the input resistance (unpaired t-test, *P* = 0.2820; control, 47.04 ± 11.70 pA; Ru360, vs. 40.87 ± 8.57 pA).

**Figure 6.**
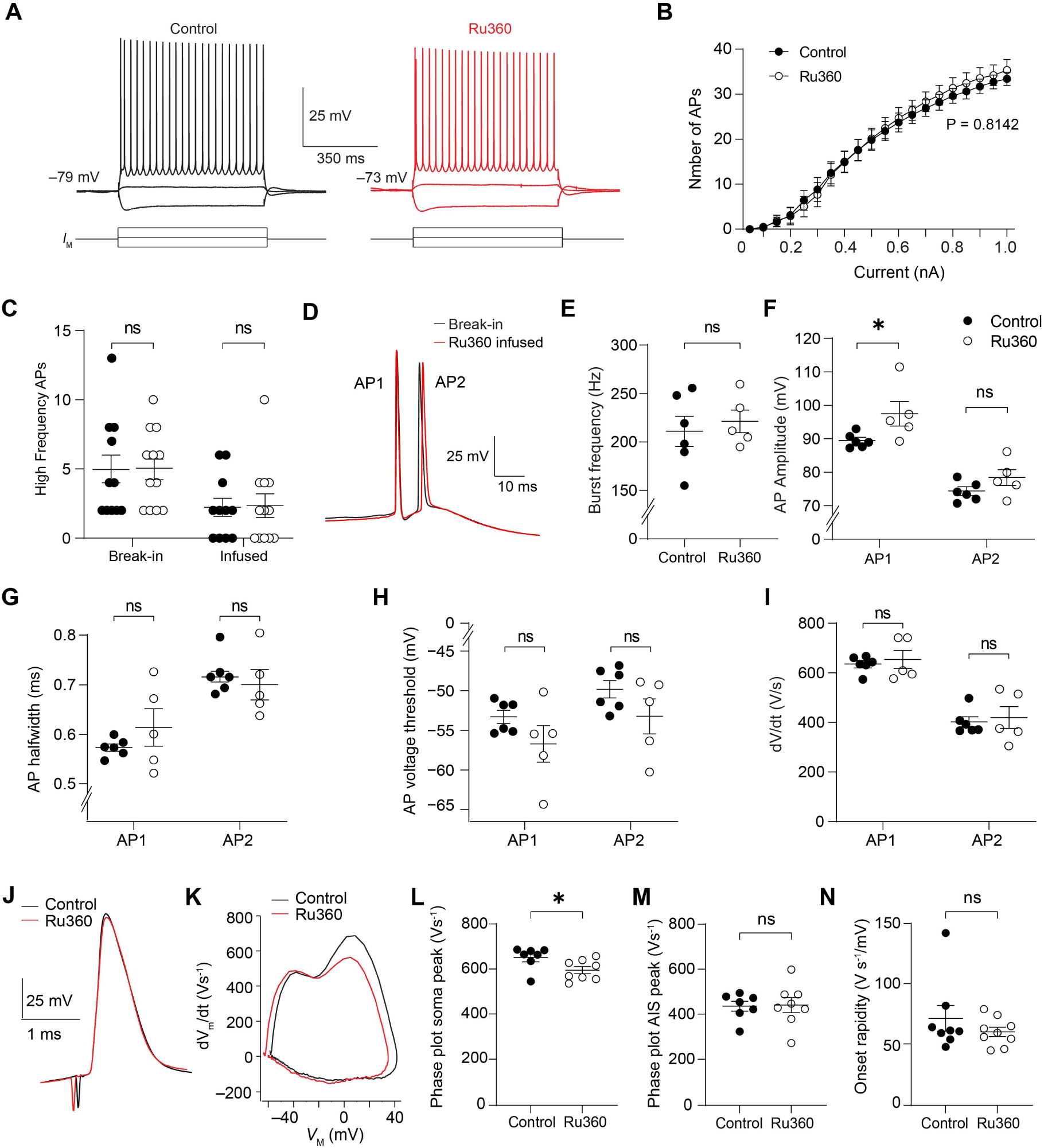
mt-Ca^2+^ does not regulate excitability, bursting or AP waveform in L5 pyramidal neurons. **A** Example traces of AP trains in a control or Ru360-infused cell; **B** Population data showing unchanged firing frequency upon somatic current injection upon Ru360 infusion (two-way ANOVA; treatment effect *P* = 0.8142, F_(1, 17)_ = 0.05041; interaction effect *P* = 0.8250, F_(19, 323)_ = 0.7046; Bonferroni *post hoc* test, control vs Ru360 *P* > 0.9999 for break-in and infused) **C** Example traces of high-frequency burst of the same cell at break-in and after Ru360 infusion. Scale bar represents 25 mV/250 ms; **D** Quantification of the number of high frequency APs at break-in or after control or Ru360 infusion (two-way ANOVA; treatment effect *P* = 0.9145, F_(1, 21)_ = 0.01181; interaction effect *P* = 0.9535, F_(1, 21)_ = 0.0003); **E** Quantification of the within-burst frequency shows no effect of mt-Ca^2+^ buffering inhibition (unpaired t-test, *P* = 0.6201); **F-I** Quantification of within-burst AP amplitude (F; two-way ANOVA; treatment effect *P* = 0.0201, F_(1, 9)_ = 7.948; interaction effect *P* = 0.3967, F_(1, 9)_ = 0.7920; Bonferroni *post hoc* test Control vs Ru360, AP1 \**P* = 0.0367, AP2 *P* = 0.4143; AP1 vs AP2 Control *P* = 0.0013, Ru360 *P* = 0.0005), half-width (G; two-way ANOVA; treatment effect *P* = 0.6952, F_(1, 9)_ = 0.1638; interaction effect *P* = 0.0076, F_(1, 9)_ = 11.72; Bonferroni *post hoc* test Control vs Ru360, AP1 *P* = 0.4582, AP2 *P* > 0.9999; AP1 vs AP2 Control and Ru360 *P* < 0.0001), voltage threshold (H; two-way ANOVA; treatment effect *P* = 0.1651, F_(1, 9)_ = 2.283; interaction effect *P* = 0.9941, F_(1, 9)_ = 5.784 × 10^−5^; Bonferroni *post hoc* test Control vs Ru360, AP1 *P* = 0.3113, AP2 *P* = 0.3092; AP1 vs AP2 Control *P* = 0.0004, Ru360 *P* = 0.0009) and d*V*/d*t* (I; two-way ANOVA; treatment effect *P* = 0.6699, F_(1, 9)_ = 0.1942; interaction effect *P* = 0.9877, F_(1, 9)_ = 0.0002518; Bonferroni *post hoc* test Control vs Ru360, AP1 and AP2 *P* > 0.9999; AP1 vs AP2 Control and Ru360 *P* < 0.0001). **J** Example traces of single APs (aligned at AP onset) elicited by a 3 ms 1nA pulse in a control and a Ru360-infused cell; **K** Example phase plane plots of the APs in J; **L-N** Population data showing unchanged rapidity of AP onset (L; Mann-Whitney test, *P* = 0.6058) and AIS component peak slope (M; unpaired t-test, *P* = 0.9064) while somatic component peak slope shows a slight decrease (N; Mann-Whitney test, *P* = 0.0289). B: control, n = 10 cells from 4 mice; Ru360, n = 10 cells from 4 mice; D: control, n = 11 cells from 6 mice; Ru360, n = 12 cells from 6 mice; E: control, n = 6 cells from 4 mice; Ru360, n = 5 cells from 4 mice; F-I: control, n = 6 cells from 4 mice; Ru360, n = 5 cells from 4 mice; L: control, n = 8 cells from 4 mice; Ru360, n = 9 cells from 3 mice; M-N: control, n = 7 cells from 3 mice; Ru360, n = 8 cells from 3 mice.

L5 pyramidal neurons are characterized by their ability to generate short high-frequency (>100 Hz) bursts, which predominantly occur during onset of membrane depolarization. Burst firing is mediated by the rapid opening of Na^+^ channels and shaped by both the Kv7 and BK channels (Battefeld *et al*., 2014; Roshchin *et al*., 2018; Niday & Bean, 2021) which are all Ca^2+^-gated (Ben-Johny *et al*., 2014; Wang *et al*., 2014; Bernardo-Seisdedos *et al*., 2018; Roshchin *et al*., 2018; Niday & Bean, 2021). We postulated that mt-Ca^2+^ regulation of membrane excitability may occur at the timescale of short inter-spike intervals and therefore its contribution could be masked when quantifying average rates. We therefore investigated the role of mt-Ca^2+^ buffering to burst properties. First, we compared the number of high-frequency APs at rheobase immediately after establishing whole-cell access with the number after 45 minutes of Ru360 or control infusion. In both the control and Ru360-infused neurons, we observed a reduction in the number of high-frequency APs, likely due to washout of the cytoplasm with the intracellular recording solution (**Figure 6C**; two-way ANOVA; infusion effect *P* < 0.0001, F_(1, 21)_ = 27.64; Bonferroni *post hoc* test, break-in vs. infusion, control *P* = 0.0028, Ru360 *P* = 0.0023; control vs Ru360 at break-in and after infusion *P* >0.9999). This reduction was however indistinguishable between the two treatments (two-way ANOVA with Bonferroni’s *post hoc* test*, P* > 0.9999). Similarly, the fraction of neurons that maintained burst firing was similar between the two groups (control, 6 out of 9 or 66.7%; Ru360, 6 out of 11 or 54.5%; *P* = 0.5231, binomial test). Secondly, in this stable group we then compared the AP properties within the burst cluster. Quantification of the burst frequency showed no difference upon mt-Ca^2+^ buffering inhibition (**Figure 6D-E**; control, 211.0 ± 38.0 Hz; Ru360, 221.3 ± 26.1 Hz; unpaired t-test). While Ru360 infusion increased the amplitude for the first AP within a burst cluster (**Figure 6F**; two-way ANOVA with Bonferroni *post hoc* test, *P* = 0.0367; control, 89.44 ± 2.21 mV; Ru360, 97.42±8.41 mV) the amplitude of the second AP was unaffected (two-way ANOVA with Bonferroni *post hoc* test, *P* > 0.9999; control, 74.48 ± 2.87 mV; Ru360, 78.20 ± 5.42 mV). Furthermore, all AP voltage thresholds, half-widths and d*V*/d*t* values were comparable between groups (**Figure 6G-I**; voltage threshold AP1 *P* = 0.3113, AP2 *P* = 0.3098; half-width AP1 *P* = 0.4582, AP2 *P* > 0.9999; d*V*/d*t* AP1 and AP2 *P* > 0.9999). Finally, we determined whether the upstroke of single APs (generated by a 3 ms pulse) is influenced by mt-Ca^2+^ buffering (**Figure 6J-N**). Upon Ru360 infusion, the AP amplitude remained constant (unpaired t-test, *P* = 0.8359), as well as the half-width (unpaired t-test, *P* = 0.5073) and the current and voltage threshold (unpaired t-tests; respectively *P* = 0.5330 and *P* = 0.7096). Phase plane plots showed a slightly reduced rate of rise peak of the somatic component of the AP (**Figure 6K-M**; Mann-Whitney test, *P* = 0.0289) without change in the AIS component (unpaired t-test, *P* = 0.9064). Similarly, the rapidness of AP onset was unchanged upon Ru360 infusion (**Figure 6N**; Mann-Whitney test, *P* = 0.6058).

In conclusion, these electrophysiological experiments indicate that mt-Ca^2+^ buffering at the AIS and soma do not influence AP generation in L5 pyramidal neurons.

## Discussion

Combining cell-type specific ultrastructural data and genetically encoded Ca^2+^ imaging tools we found that mitochondria at the AIS in neocortical L5 pyramidal neurons densely populate the proximal domain of the AIS and powerfully take up Ca^2+^. Despite the critical role of the AIS in determining AP generation the pharmacological block of mt-Ca^2+^ buffering with the selective MCU inhibitor Ru360 did, however, not impact the membrane excitability and neither showed a role in modulation of the slow AHP, AP initiation or adaptation.

The slow AHP in L5 pyramidal neurons regulates spike frequency adaptation and is detectable as an outward K^+^ current mediated by the Ca^2+^-dependent K^+^ channel isoform KCa_3.1_, lasting tens of seconds following a train of high-frequency spikes (Schwindt *et al*., 1992; Guan *et al*., 2015; Roshchin *et al*., 2020; Groten & MacVicar, 2022). A common observation across different cell types is that blocking mt-Ca^2+^ influx via the MCU prolongs the duration of cytoplasmic Ca^2+^ transients and subsequently increases the AHP duration (Groten & MacVicar, 2022; Kirchner *et al*., 2024). While our imaging of mt-Ca^2+^ in the soma and axon showed a similar time- and activity-dependent increase and a nearly complete block with Ru360 we did not find changes in the sAHP amplitude nor its time course. One possible explanation of this contradictory result could be that in adult mice, and in thick-tufted L5 pyramidal neurons, the sAHP is mediated by Ca^2+^-independent conductances. AP firing causes millimolar concentration changes in cytoplasmic Na^+^ diffusing into the distal axon and soma (Kole *et al*., 2008; Fleidervish *et al*., 2010). Electrophysiological recordings combined with Ca^2+^- and Na^+^-imaging shows that the sAHP time course also follows the internal Na^+^ concentration, is tetrodotoxin-dependent, abolished by the selective blocker ouabain and thus mediated by the outward current of the Na^+^/K^+^-ATPase pump (Gulledge *et al*., 2013). A contribution of cytoplasmic Ca^2+^ entry to the sAHP was only visible when reducing the temperature to ∼23 °C. In addition to temperature, differences between the relative contributions of Ca^2+^ dependent K^+^ channels and the Na^+^/K^+^-ATPase pump to the sAHP include the subtype of L5 neurons (thick-versus slender-tufted) and age (Gulledge *et al*., 2013; Guan *et al*., 2015; Tiwari *et al*., 2018). Although mt-Ca^2+^ influx is also required for oxidative phosphorylation and thereby expected to influence the Na^+^/K^+^-ATPase pump current, recent studies indicate intramitochondrial Ca^2+^ determines only ∼15% of ATP while the majority is produced by alternative, extramitochondrial, Ca^2+^ handling mechanisms (Nichols *et al*., 2016; Szibor *et al*., 2020). Another possible explanation may be that mitochondria are molecularly distinct between neuronal subtypes, resulting in variable Ca^2+^ buffering capacities (Fecher *et al*., 2019).

### AP initiation and burst firing is independent from mitochondrial Ca^2+^ buffering

While the currents for the slow AHP in thick-tufted L5 pyramidal neurons may be independent of mitochondria Ca^2+^ uptake, voltage-gated Ca^2+^ influx and cytoplasmic Ca^2+^ levels are well established sources to regulate AP repolarization and width, as well as the inter-spike intervals (Schwindt *et al*., 1992; Bender & Trussell, 2009; Gründemann & Clark, 2015; Hanemaaijer *et al*., 2020; Filipis *et al*., 2023). Analysis of somatically recorded APs, which reflect both AIS and somatodendritic excitability, revealed however no coherent evidence that the kinetics and threshold properties of the AP were changed upon MCU block. The lack of a role of mt-Ca^2+^ on BK channel-mediated excitability is not unexpected. Ca^2+^ activated BK channels, regulating the fAHP within milliseconds possibly via Ca^2+^ domains near the plasma membrane, possess sub-millisecond activation kinetics (Niday & Bean, 2021; Filipis *et al*., 2023). However, AIS mt-Ca^2+^ buffering follows cyt-Ca^2+^ transients with a ∼150 ms temporal delay, in good agreement with previous reports (Ashrafi *et al*., 2020; Groten & MacVicar, 2022; Stoler *et al*., 2022). In contrast, Ca^2+^ -gated Kv7 channels are open at rest and require tens of milliseconds to fully activate and regulate spike-frequency adaptation (Brown & Passmore, 2009; Battefeld *et al*., 2014). However, since neither steady-state AP firing nor the rheobase were affected, it is unlikely Kv7 channels are modified in the absence of mt-Ca^2+^ uptake (**Figure 5**). Together, based on various lines of electrophysiological analyses, our data indicate that mt-Ca^2+^ sequestration buffering does not play a role in the baseline AP properties of L5 pyramidal neurons.

### Technical limitations

In the present study, we acutely blocked mt-Ca^2+^ buffering in L5 pyramidal neurons by intracellular infusion of Ru360 in the whole-cell configuration. Although this mode enables high temporal resolution electrophysiological recordings with low access resistance (< 20 MW) and previously was used to show a role of the MCU to neuronal excitability (Groten & MacVicar, 2022; Kirchner *et al*., 2024), it comes with the disadvantage of altering the intracellular ionic composition and/or phosphorylation, potentially masking a mt-Ca^2+^ contribution to membrane properties. An alternative not perturbing the intracellular environment would be to use perforated patch-clamp recordings. While this prohibits Ru360 infusion an interesting alternative for future studies would be to combine perforated patch-clamp recordings with optical approaches to locally disrupt AIS mt-Ca^2+^ buffering (Tkatch *et al*., 2017; Tjiang & Zempel, 2022). Although currently available tools do not specifically target mt-Ca^2+^ uptake, an exciting possibility could be to develop light-mediated disruption of *e.g.* the MCU (Hermann *et al*., 2015). Another possible limitation of the present study is that mitochondria are not the only Ca^2+^ buffering organelle at the AIS. Selectively in the thick-tufted L5 pyramidal neurons of the primary somatosensory cortex, the AIS harbours a giant saccular organelle, a large variant of the cisternal organelle consisting of smooth endoplasmic reticulum (ER), which determines by Ca^2+^-induced Ca^2+^ release ∼50% of the AP-dependent Ca^2+^ rise (Antón-Fernández *et al*., 2015; Hanemaaijer *et al*., 2020). Although genetic ablation of the cisternal organelle does not impact AP firing rates (Orth *et al*., 2007), it remains to be tested whether mitochondrial and ER Ca^2+^ handling interact to shape cytoplasmic Ca^2+^ homeostasis at the AIS.

### Anatomical distribution and non-electrical roles of AIS mitochondria

The present finding that AP initiation and spike adaptation were independent from the MCU-associated Ca^2+^ uptake is consistent with the generally preserved cognitive and motor control under baseline conditions in the MCU knockout mice (Pan *et al*., 2013; Nichols *et al*., 2016; Szibor *et al*., 2020). The substantially lower number of mitochondria in the AIS relative to the soma (**Figures 1, 2**) is in good support of recent emerging insights of their distribution in mammalian motor neurons, human induced pluripotent stem cells and drosophila AIS (Tamada *et al*., 2021; Tjiang & Zempel, 2022; Wodrich *et al*., 2024). Here, we extend these observations by identifying proximal-distal gradients of mitochondria, with significantly smaller ones in the distal domain of the AIS. The molecular mechanisms determining the mitochondria distribution along the AIS subdomains are not well understood. One candidate is Ca^2+^ dependent anchoring of mitochondria by for instance the adaptor protein Miro or the anchoring protein syntaphilin (Devine & Kittler, 2018). However, this possibility is not very likely since Ca^2+^ influx is uniformly high along the AIS and mitochondrial clustering also occurs in the absence of Ca^2+^ influx, as shown in basket cell internodes (Kole *et al*., 2022). Another possibility is that mitochondria are tethered to the endoplasmic reticulum, for instance via vesicle-associated membrane protein-associated proteins (Bapat *et al*., 2024) or the axonal membrane (Lackner *et al*., 2013). To our knowledge, mitochondria are not physically tethered to voltage-gated ion channels but given their spatially nonuniform distributions along the proximal-distal axis (Kole & Stuart, 2012; Jenkins & Bender, 2024) it is interesting to speculate that mechanisms defining subdomain localization may be (partially) shared or that the ion channel distribution guides mitochondrial clustering.

Since AP initiation typically emerges from the distal AIS region (Kole & Stuart, 2012; Jenkins & Bender, 2024), our anatomical and functional data strongly suggest that under baseline conditions mitochondrial Ca^2+^ handling is dispensable for AP initiation. If excitability is not affected what could be the role of mitochondria in the AIS? One intriguing possibility is that the MCU-mediated Ca^2+^ importing at the AIS may be involved during prolonged periods of elevated cytoplasmic Ca^2+^ and act as instructive for cellular homeostasis and structural plasticity, analogous to their role at spines and presynaptic terminals (Kwon *et al*., 2016; Lewis *et al*., 2018). In line of this idea, global MCU knockout preserves baseline activity levels and mitochondrial ultrastructure but renders cortical neurons vulnerable as the respiratory capacity fails to offset the increased glycolytic demands (Nichols *et al*., 2016; Szibor *et al*., 2020). For example, during strongly increased levels of neuronal activity, the distal AIS shortens via endocytosis of voltage-gated Na^+^ channels and Ankyrin G proteins from the distal AIS plasma membrane within a period of tens of minutes (Jamann *et al*., 2021; Fréal *et al*., 2023). AIS plasticity induced by neuronal activity or brief N-methyl-D-aspartate receptor activation is a Ca^2+^-dependent process (Evans *et al*., 2013; Fréal *et al*., 2023). More extreme AIS plasticity induction protocols found that AIS disassembly requires proteasomes. It is likely that mitochondria dock to sites of the AIS cytoskeleton during remodelling. An interesting experiment would be to image mt-GFP during AIS plasticity and genetically silence the MCU.

More evidence for a functional role of mitochondria is during axonal damage such as demyelination and/or neurodegenerative diseases when the organelles increase in number and size within the AIS, even accumulating at the distal AIS (Sasaki & Iwata, 1996; Tamada *et al*., 2021; Wodrich *et al*., 2024). In demyelinated axons the mitochondrial anchoring protein syntaphilin is required for the immobilization, clustering and the increase of mitochondria size (Ohno *et al*., 2014). AIS mitochondria under pathological conditions may support the adaptation of neuronal excitability or axonal cargo transport. For example, motor neurons in amyotrophic lateral sclerosis (ALS) accumulate mitochondria at the AIS, are characterized by a shorter AIS length and hypoexcitability (Sasaki & Iwata, 1996; Harley *et al*., 2023). Using a nerve injury model combined with ALS genetic engineering proteasome-associated disassembly AIS was found to be required to enable mitochondrial transport beyond the AIS and support injury repair (Kiryu-Seo *et al*., 2022). The complex and bidirectional interactions between AIS integrity, mitochondria and injury remain to be further studied. Interestingly, pharmacologically enhancing mt-Ca^2+^ buffering in hippocampal neurons rescues epilepsy symptoms, in which mt-Ca^2+^ buffering at the AIS could be involved (Styr *et al*., 2019). Taken together, the current findings show that L5 pyramidal neurons regulate their axonal mitochondrial content at the microdomain level, that mitochondria cluster to the proximal AIS but are dispensable for maintaining normal levels of membrane excitability. These data open avenues to explore the roles of mitochondria in the (dis)assembly and maintenance of the AIS during plasticity and pathology.

## Acknowledgements

This study was funded by a ZonMW Off Road grant 04510012010066 to K.K. and a Netherlands Research Council (NWO) Vici grant 865.17.003 to M.K. The authors are indebted to the members of the Axonal Signaling department of the NIN for discussing preliminary data and manuscript versions, and Naomi Petersen for the support with AAV injections.

## Methods

### Mice

All animal procedures were performed after evaluation by the Royal Netherlands Academy of Arts and Sciences (KNAW) Animal Ethics Committee (DEC) and Centrale Commissie Dierproeven (CCD, license AVD8010020172426). The specific experimental designs with animals were evaluated and monitored by the Animal Welfare Body (IvD, protocols NIN19.21.09, NIN19.21.12 and NIN21.2108). The mouse strain used for this research project was B6.FVB(Cg)-Tg(Rbp4-cre)KL100Gsat/Mmucd, RRID:MMRRC_037128-UCD, obtained from the Mutant Mouse Resource and Research Center (MMRRC) at University of California at Davis, an NIH-funded strain repository, and was donated to the MMRRC by MMRRC at University of California, Davis. Mice, here called *Rbp4-Cre*, were made from the original strain MMRRC:032115 donated by Nathaniel Heintz, Ph.D., The Rockefeller University, GENSAT and Charles Gerfen, Ph.D., National Institutes of Health, National Institute of Mental Health. Mice (8–11 wks old) were kept at a 12 h day-night cycle with ad libitum access to food pellets and water. Cages were open or IVC cages with corncob bedding. Male or female *Rbp4-Cre* mice were housed together with at least one cage mate. Ambient temperature was maintained at 20–24 °C, humidity at 45– 65%.

### AAV production

Plasmids to generate mt-GFP or mt-GCaMP6f AAVs were generated as described previously (Kole *et al*., 2022). For AAV production, HEK293-T cells of low (<25) passage were maintained in Dulbecco’s modified Eagle’s medium (DMEM, Thermo Fisher Scientific #31966-047). The medium contained 10% foetal calf serum (FCS, Thermo Fisher Scientific A4766801) and 5% penicillin-streptomycin (Pen-Strep, Thermo Fisher Scientific #15140122), cells were maintained at 37 °C and 5% CO_2_. For virus production, cells were seeded in 15 cm dishes at a density of 1–1.25 × 10^7^ cells per dish. The following day, 1–2 h before transfection, the DMEM was replaced by fresh Iscove’s modified Dulbecco’s medium (IMDM, Sigma Aldrich I-3390) containing 10% FCS, 5% Pen-Strep and 5% glutamine (Thermo Fisher Scientific #25030081). For transfection, AAV rep/cap, AAV helper (mt-GFP: AAV5; mt-GCaMP6f: AAV1) and transfer plasmids were mixed and diluted in saline before mixing with saline-diluted polyethylenimine (PEI, Polysciences #23966-2) and brief vortexing. After incubation (20–25 minutes), the transfection mix was added to the culture plates in a drop-wise fashion. After 16 hours, the medium was refreshed after which the cells were left for an additional 72 hours. Then, medium was discarded, and cells were then collected. Lysis to release AAVs was achieved via three freeze-thaw cycles. Cell lysate was then loaded on an iodixanol gradient (60, 40, 25 and 15% iodixanol, ELITechGroup #1114542) in Beckman Quick-Seal Polyallomer tubes (Beckman-Coulter #342414). Centrifugation was performed at 16 °C and 69,000 rpm (488727.6 × *g*) for 1 h and 10 min in a Beckman-Coulter Optima XE-90 Ultracentrifuge using a Type 70 Ti rotor.

The virus-containing fraction was then extracted from the tubes and AAVs were then concentrated in Dulbecco’s Phosphate Buffered Saline (D-PBS) + 5% sucrose using Amicon Ultra-15 (100 K) filter units (Merck Millipore UFC910024) at 3220 *× g*. To ensure complete replacement of iodixanol with D-PBS + 5% sucrose, at least 4 rounds of centrifugation were used. Viral titres were determined using quantitative PCR (titres: mt-GFP-DIO, 4.43 × 10^13^ gc/ml; mt-GCaMP6f-DIO, 3.44 × 10^13^ gc/ml). Viral aliquots were stored at −80 °C until further use.

qPCR primers, recognize AAV2 inverted terminal repeats (ITRs): 5′-GGAACCCCTAGTGATGGAGTT-3′ 5′-CGGCCTCAGTGAGCGA-3′

### Viral injection

Viral injections were typically performed at 8–9 weeks of age. Mice were anaesthetized using isoflurane (induction, 3%; maintenance, 1.2–1.5%) after which they received 5 mg/kg Metacam subcutaneously. Using a heating pad, the body temperature was monitored and maintained at 37 °C. To prevent the eyes from drying out, eye ointment was applied. Then, the head was shaved using electronic clippers and hair removal cream, after which it was placed in a stereotaxic frame (Kopf). An incision was made in the skin along the midline. Before removing the periost, lidocaine (10%) was administered locally. Small (<1 mm) bilateral craniotomies were made at −0.5 mm caudally from Bregma and 2.5 mm laterally from the midline. Care was taken not to damage the dura mater. Using a Nanoject III (Drummond) fitted with a sharp glass pipette, 40–50 nL of virus was injected at 1 nL/s and at a depth of 450 µm. Approximately 3 min after finishing the injection, the needle was retracted slowly. Bone wax was applied to the craniotomies and the skin was carefully sutured before mice were allowed to recover. During the 3–5 days following surgery, mice were monitored closely. Their weight, locomotion and overall wellbeing were checked.

### Acute slice preparation

After 2–3 weeks of virus expression, mice were deeply anaesthetized using pentobarbital (50 mg/kg, intraperitoneal injection) and transcranially perfused with ice-cold, carbogenated (95% O_2_, 5% CO_2_) cutting artificial cerebrospinal fluid (cACSF; 125 mM NaCl, 3 mM KCl, 6 mM MgCl_2_, 1 mM CaCl_2_, 25 mM glucose, 1.25 mM NaH_2_PO_4_, 1 mM kynurenic acid and 25 mM NaHCO_3_). The brain was quickly dissected out, after which 400 µm thick parasagittal slices were cut using a Vibratome (1200S, Leica Microsystems) all the while keeping the brain submerged in ice-cold carbogenated cACSF. Slices were immediately transferred to a holding chamber containing carbogenated cACSF at 35 °C where they were kept for 35 min to recover. After this period, they were allowed to return to room temperature for at least 30 min before starting experiments.

### Electrophysiology and two-photon imaging

Slices were transferred to a recording chamber with continuous perfusion (1–2 ml per minute) of carbogenated recording ACSF (rACSF, 125 mM NaCl, 3 mM KCl, 1 mM MgCl_2_, 2 mM CaCl_2_, 10 mM glucose, 5 mM L-Lactate, 1.25 mM NaH_2_PO_4_, and 25 mM NaHCO_3_). Bath temperature was maintained at 32 °C. Glass pipettes with an open tip impedance of 6–7 MΩ were filled with an intracellular solution containing (130 mM K-Gluconate, 10 mM KCl, 10 mM HEPES, 4 mM Mg-ATP, 0.3 mM Na_2_-GTP, 10 mM Na_2_-phosphocreatine; pH ∼7.25, osmolality ∼280 mOsmol/kg). In a subset of experiments, the intracellular solution was supplemented with 5 mg/ml biocytin (Sigma-Aldrich, B4261) and 50 µM Atto-594 (Sigma-Aldrich, A08637). Whole-cell recordings were made using a patch-clamp amplifier (Multiclamp 700B, Axon Instruments, Molecular Devices, RRID: SCR_018455) operated by AxoGraph X software (version 1.5.4; RRID: SCR_014284). Action potential trains were evoked using 700 ms step pulses, from −250 pA to +1000 nA with increments of 50 pA. Starting below spike threshold, 3 ms incremental (2.5–5 pA) step pulses were used to evoke single APs. Voltage was digitally sampled at 100 Hz using an AD/DA converter (ITC-18, HEKA Elektronik GmbH). During current-clamp experiments, the access resistance (range: 15–30 MΩ) was fully compensated using bridge balance and capacitance neutralization of the amplifier. Membrane potentials in this study are corrected for a –14 mV junction potential of the intracellular recording solution. Somatic single-cell recordings were made from mt-GFP^+^ or mt-GCaMP6^+^ cells, which were visualized using a two-photon (2P) laser-scanning microscope (Femto3D-RC, Femtonics Inc., Budapest, Hungary). Imaging was controlled using MES software (Femtonics Inc., Budapest, Hungary, version 6.3.7902). To visualize mt-GFP/mt-GCaMP6f, a Ti:Sapphire pulsed laser (Chameleon Ultra II, Coherent Inc., Santa Clara, CA, USA) was tuned to 770 nm for two-photon excitation. Fluorescent signals were detected using two photomultipliers (PMTs, Hamamatsu Photonics Co., Hamamatsu, Japan), one for mt-GFP and for Cal590. For motility imaging, *z*-stacks were acquired every 10 s for 8–12 min. The Image Stabilizer plugin for FIJI was used for drift correction. For Ca^2+^ imaging, mt-GCaMP6f^+^ cells were targeted for single-cell patch clamping and filled with Cal-590 (20 µM; Sigma 08637). Next, *z*-stacks were made with the laser tuned to 800 nm to identify the axon and its subcompartments (AIS, myelinated internode and nodes). Mt- and cyt-Ca^2+^ responses were then simultaneously visualized with the laser tuned to 940 nm and recorded at ∼20 Hz imaging frequency. Optical and electrophysiological recordings were synchronized via a TTL pulse from the microscope to the amplifier. Ca^2+^ responses were analysed using a custom-written Matlab script (Kole *et al*., 2022). Briefly, regions of interest were selected from which background was subtracted, bleach correction and smoothing were applied, and ΔF/F was calculated.

### Immunohistochemistry

For staining of biocytin-filled cells, upon completion of experiments acute slices were immediately placed into 4% PFA in PBS for 20–25 min, followed by 3 washes of 10 min with PBS. Next, blocking was performed in PBS containing 1–2% Triton-X and 10% normal goat serum for 2 h. After blocking, sections were moved to blocking buffer containing primary antibodies (see **Supplemental Table 1**) and were incubated overnight at room temperature. Following three 10 min washes in PBS, sections were transferred to PBS containing secondary antibodies and were incubated overnight 2 h room temperature or overnight at 4 °C. The tissue was finally washed again in PBS three times for 10 min before mounting using FluorSave mounting medium (Merck-Millipore #345789). For immunostainings against MBP, the staining protocol was adjusted: blocking was done 1 h at 37 °C and 1 h room temperature and using 1% Triton, tissue was washed only once in PBS per washing step, and secondary antibodies were incubated 1 h at 37 °C and 1 h room temperature. For immunostainings on sections without biocytin-filled cells, 400 µm PFA-fixed sections were first cryoprotected by placing them into 30% sucrose-PBS solution until fully saturated. A sliding freezing microtome (Zeiss Hyrax S30; temperature controlled by a Slee Medical GmbH MTR fast cooling unit) was used to cut 40 µm sections, which were either placed in PBS for immediate use or stored at −20 °C in cryoprotectant solution (30% ethylene glycol, 20% glycerol, 0.05 M phosphate buffer) until further use. Immunostaining protocol was the same as for 400 µm sections, but the blocking buffer contained 0.5% Triton-X and incubation duration for secondary antibodies was 2 h at room temperature. All steps are performed with gentle shaking, except for the 37 °C incubation steps. For analysis of normal and demyelinated subcortical white matter, *Rbp4-Cre* mice injected with AAV1-EF1a-mCherry-DIO and AAV5-EF1a-mtGFP-DIO (mixed 1:1) were first perfused with 1x PBS followed by 4% PFA-PBS. Brains were then dissected out and allowed to fix O/N in 4% PFA-PBS after the brains were cryoprotected using 30% sucrose-PBS and processed into 40 µm coronal sections as described above.

### Confocal microscopy

For confocal imaging, a Leica SP8 X confocal laser-scanning microscope controlled by Leica Application Suite AF (version 3.5.7.23225) was used. Biocytin-filled cells were imaged with a 63× oil-immersion lens. The tile-scan function was used with automated sequential acquisition of multiple channels enabled, and step sizes in the Z-axis were 0.3–0.75 µm, and images were collected at a 2048 × 2048 pixel resolution at 100– 150 Hz. Axonal reconstructions and quantification of mitochondrial density were performed manually using Neurolucida Software (MBF Bioscience, version 2019.2.1 or 2020.1.3, 64 bit, RRID: SCR_001775). Care was taken to include only mitochondria of which the mt-GFP signal was located clearly inside the cytosol and followed the path of the reconstructed neurite. Upon completion of tracing, analysis of reconstructions was performed using Neurolucida Explorer (MBF Bioscience, 2019.2.1, RRID: SCR_017348). FIJI (FIJI 64 bit; ImageJ version 1.53q; RRID: SCR_002285) was used to extract partial images for use in figures and generation of the kymograph in Figure 3. Motile mitochondria were identified by eye, mitochondria were considered stable if they displaced less than 2 µm.

### 3D Electron microscope data analysis

3D EM data was obtained from the Microns dataset (Consortium *et al*., 2025) (www.microns-explorer.org). This dataset consists of a 1 mm^3^ EM block of primary visual cortex of one P87 male mouse expressing GCaMP6s in excitatory neurons (*Slc17a7*-Cre and Ai162 heterozygous transgenic lines; JAX stock 023527 and 031562, respectively). L5 pyramidal neurons were identified based on their distance from the pia as well as their morphology (*i.e.* their pyramidal shaped soma, large apical dendrite and axon projecting into the white matter). EM volumes containing the AIS and the first internode, as well as their cytosolic segmentation were then downloaded, after which the mitochondria were segmented and the segment length traced manually using Volume Annotation and Segmentation Tool (VAST version 1.4.1) (Berger *et al*., 2018). VAST Tools Matlab scripts were used for length and volume measurements and exports. Exported 3D models were then rendered using 3ds Max (Autodesk, version 25.0.0.997, SCR_014251). See **Supplementary Table 2** for cells used in the analysis.

### Statistics and reproducibility

Prism (Graphpad, version 8.4.3, RRID: SCR_002798) was used for all statistical comparisons. Dataset normality was determined using D’Agostino & Pearson or Shapiro-Wilk tests. If data deviated significantly from a normal distribution we used non-parametric tests. For comparisons between groups we used a two-tailed unpaired t-test (normal data) or a two-tailed Mann–Whitney test (non-normal data). To test interactions between groups and treatments two-way ANOVA was used; Bonferroni’s post hoc test (normal data) or Dunn’s post hoc test (non-normal data) was used for multiple comparisons. To avoid overpowering of non-nested statistical tests, a nested t-test was used when large numbers of datapoints were involved (*i.e.* mitochondrial contours). Figure legends contain *P*-values and *n*-numbers and whether the latter signify mice, cells or mitochondria. Means are presented with SEM. To ensure reproducibility, data was collected from multiple cells from multiple mice: experiments were replicated in at least 4 cells from 3 mice. The only exception is the 3D EM reconstructions which were done using data obtained from one mouse. Cells were randomly infused with either control or Ru360-containing intracellular solution. Electrophysiological data was excluded if cells had an unstable resting membrane potential between sweeps.

---

**Supplementary Table 1.**
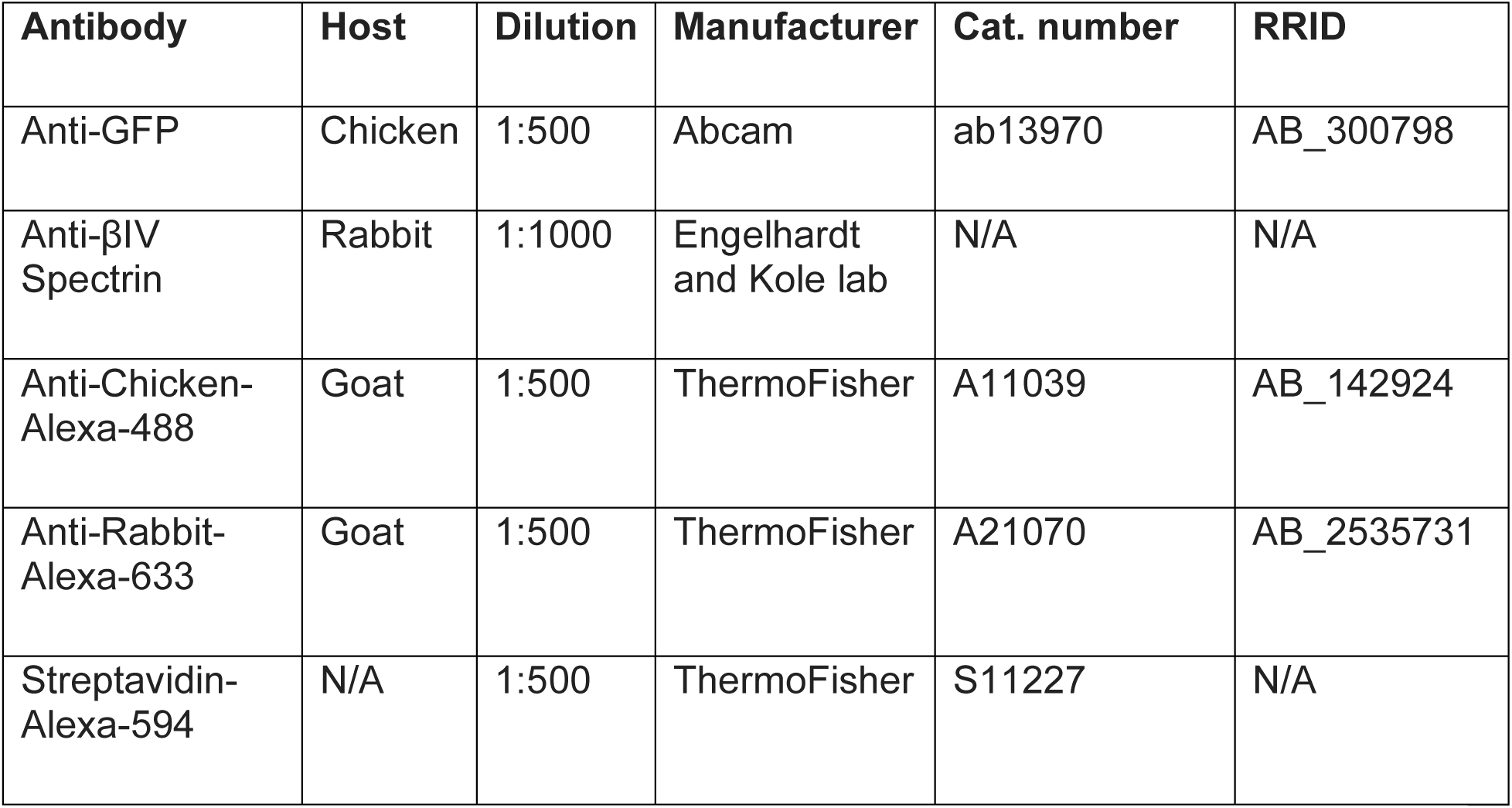
Overview of antibodies and dilutions used in this study.

| **Antibody** | **Host** | **Dilution** | **Manufacturer** | **Cat. number** | **RRID** |
| --- | --- | --- | --- | --- | --- |
| Anti-GFP | Chicken | 1:500 | Abcam | ab13970 | AB_300798 |
| Anti-βIV Spectrin | Rabbit | 1:1000 | Engelhardt and Kole lab | N/A | N/A |
| Anti-Chicken-Alexa-488 | Goat | 1:500 | ThermoFisher | A11039 | AB_142924 |
| Anti-Rabbit-Alexa-633 | Goat | 1:500 | ThermoFisher | A21070 | AB_2535731 |
| Streptavidin-Alexa-594 | N/A | 1:500 | ThermoFisher | S11227 | N/A |

**Supplementary Table 2.** Overview of the cells used in the 3D EM analysis. Cell L5PN_7 was used as example in Figure 1.

| **Cell name** | **Neuroglancer ID** | **Soma coordinates** |
| --- | --- | --- |
| L5PN_1 | 864691135280056225 | 181036, 189486, 20898 |
| L5PN_2 | 864691135463505821 | 204687, 202669, 20853 |
| L5PN_3 | 864691135293275574 | 201708, 198355, 21239 |
| L5PN_4 | 864691136990817685 | 227311, 197012, 21697 |
| L5PN_5 | 864691135384023787 | 209082, 201077, 21549 |
| L5PN_6 | 864691135725697451 | 219205, 190682, 22103 |
| L5PN_7 | 864691135408575689 | 214530, 192267, 22033 |

